# A gene-agnostic FACS technique to isolate stem cells in *Hydractinia symbiolongicarpus* validated with cytology

**DOI:** 10.64898/2026.07.22.740159

**Authors:** Zachary M. Lane, Christine E. Schnitzler

## Abstract

*Hydractinia symbiolongicarpus* is a powerful model for stem cell research and maintains a population of pluripotent adult stem cells throughout its lifetime. Here we describe a gene expression-agnostic FACS technique to isolate a live cell population from *Hydractinia* feeding polyps that appear to be stem cells. This technique utilizes only the general cellular component stains DAPI, DRAQ5, Calcein AM, and Pyronin Y. The stem cell population was identified via subtractive gating based on samples whose stem cell populations had been selectively depleted with the DNA-alkylating agent Mitomycin C. To validate the identity of the isolated population, a colorimetric cytological assay capable of simultaneously discriminating between all major *Hydractinia* cell types in a live-dissociated cell solution was developed using May-Grünwald and Giemsa stains. The isolated cell population was significantly depleted by Mitomycin C administration, had a high RNA content, was proliferative, had a cytological profile that matched that of Piwi1^+^ stem cells, and was ∼10x enriched with Piwi1^+^ stem cells compared to whole cell suspension, all of which support the conclusion that the isolated population is indeed comprised of stem cells. This gene-agnostic FACS technique will serve future research into *Hydractinia* stem cell biology by enabling the use of isolated populations of live stem cells in transplantation, cell culture, and spheroid experimentation, and may serve as a reference for the development of new methods in other cnidarian species.

## INTRODUCTION

*Hydractinia symbiolongicarpus* is colonial marine hydrozoan with whole-body regenerative capabilities which are orchestrated by a population of adult pluripotent stem cells (aPSCs) that it maintains throughout its lifetime (Müller et al., 1986, 2004; Varley et al., 2023). In addition to its capacity for whole-body regeneration, *Hydractinia* aPSCs are thought to contribute to this species’ lack of senescence (Gahan et al., 2016; Martínez, 1998), propensity for wound healing (Bradshaw et al., 2015; Salinas-Saavedra et al., 2023), ability to reaggregate/regenerate following complete cellular dissociation (Müller et al., 1986; Schmid et al., 1981), and natural capacity to form spheroids from dissociated cell suspensions (Curantz et al., 2025). In the laboratory, *Hydractinia* maintains stable transgenesis (Chrysostomou et al., 2022; DuBuc et al., 2020; Varley et al., 2023), can be asexually propagated via explantation indefinitely (Frank et al., 2020), can be sexually crossed with high levels of control (Cadavid et al., 2004), and readily forms chimeras and accepts tissue graphs from others of its species (Müller, 1982; Müller et al., 2004; Varley et al., 2023). *Hydractinia* husbandry is straightforward (Frank et al., 2020), and its amenability to experimental manipulation make it a powerful model for investigating aPSC maintenance and control during homeostasis, regeneration, wound healing, and aging.

The primary goal of the current study was to develop a fluorescence activated cell sorting (FACS) technique for selectively isolating live *Hydractinia* aPSCs for use in various downstream applications which require isolation of living cells such as experimental transplantation, cell culture, or spheroid generation. To accomplish this goal, we pioneered an expression-agnostic FACS approach (i.e., an approach using only general cellular component stains), similar to that established for planarian aPSC isolation (Hayashi et al., 2006; Molinaro et al., 2021). In this approach, we used the DNA-alkylating agent Mitomycin C (MitC) to selectively deplete the aPSC population of intact *Hydractinia* colonies (Müller, 1968; Müller et al., 2004) which, in turn, allowed for these cells to be identified in untreated animals via subtractive flow cytometry analysis (Hayashi et al., 2006; Molinaro et al., 2021).

Previously, flow cytometry analyses of living *Hydractinia* aPSCs have been conducted using Piwi1 transgenic reporter lines designed to fluorescently label aPSCs (Chrysostomou et al., 2022; DuBuc et al., 2020; Varley et al., 2023). This approach was specifically avoided here for the following reasons: (1) in *Hydractinia*, as in other species, expression of Piwi proteins within stem cell populations is inconsistent, and Piwi1 expression-based approaches likely isolate only a portion of the total aPSC population (Quiroga-Artigas et al., 2022; Song et al., 2025; Waletich et al., 2024); (2) Piwi1-based transgenic animals show fluorescence in non-stem cells due to both native Piwi expression in those cells (e.g., high expression in gametes, ubiquitous low expression in many cell types) (Song et al., 2025; Waletich et al., 2024) and the potential of fluorescent proteins to persist in progeny following differentiation (Corish and Tyler-Smith, 1999); (3) the development of a cell isolation technique that depends on a specific transgenesis limits potential downstream applications by requiring them to be compatible with the transgenes used in sorting (e.g., a Piwi1 transgenic based sorting approach would not allow the isolation of aPSCs from other transgenic lines of interest and all downstream experiments would have to be compatible with Piwi1-associated fluorescence, regardless of whether or not that fluorescence profile was optimal), and (4) by utilizing an approach based on general expectations of stem cell biology (e.g., high RNA content, sensitivity to cytostatic drugs), rather than specific gene expression, we aim to provide a technique with wide applications and accessibility (Müller, 1968; Rhee and Bao, 2009; Rumman et al., 2015; Van Zant, 1984). In short, our expression-agnostic approach was chosen to ensure its wide utility and accessibility, and to avoid potential biases or inaccuracies that might be introduced from using a gene-specific approach to isolate a cell type with highly variable expression (Song et al., 2025; Waletich et al., 2024).

To confirm that our FACS technique effectively isolated *Hydractinia* aPSCs, we developed a cytological staining assay that allows cell type to be assigned to live-dissociated *Hydractinia* cells based on staining patterns and diameter measurements. This was necessary because following enzymatic dissociation, live dissociated cells become spherical, losing the distinct, cell type-specific morphology typically used to discriminate between different cell populations in macerated (i.e., dissociated from fixed tissues) cell suspension in cnidarians (David, 1973; Schmid et al., 1981), thus resulting in a suspension of cells that cannot be accurately differentiated based on morphology alone. We had reason to believe that such an assay could be successfully developed thanks to previous work in *Hydractinia* which has shown the effectiveness of histological staining in cell type discrimination within whole mounts and tissue sections (Müller, 1964, 1967; Müller et al., 2004; Weismann, 1883), and the use of similar assays to discriminate between cell types in suspension for other organisms (e.g., human blood cells). We chose to utilize a general cytological assay, rather than cell type-specific expression-based labeling, to ultimately validate our FACS technique due to the inconsistent expression of marker genes in *Hydractinia* aPSCs previously mentioned and the potential for a cytological assay to enable simultaneous discrimination of all cell types without *a priori* expectations of suspension composition or the spectral limitations of fluorescence multiplexing required for simultaneous expression-based labeling. We verified our cytological cell type identification method with Piwi1 antibody staining for aPSCs and hybridization chain reaction fluorescent *in situ* hybridization (HCR-FISH) with a set of previously validated cell type marker genes for 10 additional cell types (Song et al. 2025).

## MATERIALS AND METHODS

### Animal Husbandry

Adult *Hydractinia symbiolongicarpus* colonies (291 − 10, male) were maintained at the University of Florida, Whitney Laboratory for Marine Bioscience. Colonies were grown on glass microscope slides and cultured in 38 L tanks filled with artificial seawater (Coral Pro Salt, Red Sea) at 30 ppt and kept at 18–20 °C under a 10 h/14 h light/dark regime. Animals were fed five times a week with 3-day-old SEP-Art *Artemia* nauplii (INVE Aquaculture), which were enriched two times with S.presso (SELCO) the day before colony feeding.

### Preparation of Live Cell Suspension

Feeding polyps were dissected from the colony at their base in filtered (0.22 μm) artificial seawater at 30 ppt. Dissected feeding polyps were then gently rinsed three times in 30 ppt calcium and magnesium-free artificial seawater (CMFASW) with 0.1% EGTA (2.5 mM) and placed in 75 U ml^-1^ Pronase E (Santa Cruz Biotechnology) solution in 30 ppt CMFASW with 0.1% EGTA. Feeding polyps were incubated in the Pronase solution at room temperature until a single-cell suspension was achieved, typically 1.5-2.5 hours, and were gently resuspended via pipetting every 20 minutes throughout this incubation. Once a single-cell suspension was achieved, the suspension was run through a 70 μm net filter (pluriSelect USA mini-strainer) into a low-bind microcentrifuge tube (Eppendorf DNA LoBind Tube 2.0 ml), pelleted at 500 rcf for 5 minutes at 4°C and resuspended in ice-cold 30 ppt CMFASW three times, and then filtered a final time through a 40 μm net filter (Fig. S1).

### Sequential Staining of Cell Suspension for Cytological Assay Development

Following dissociation, cells were suspended in 500 µl of ice-cold 30 ppt CMFASW and then 500 µl of ice-cold, modified ACME solution (García-Castro et al., 2021) was added dropwise, stirring gently between drops, to a final volume of 1 ml. The modified ACME solution was comprised of 60% 50 ppt CMFASW, 20% methanol, 10% glacial acetic acid, and 10% glycerin yielding a final solution with a salinity of 30 ppt that served as a gentle, isotonic fixative. The cells were then incubated in the fixative for 2 hours at room temperature on a rocker. Following the incubation, the fixed cells were pipetted onto Superfrost Plus microscope slides (Fisher Scientific) inside of circular wells drawn with an ImmEdge® hydrophobic barrier PAP pen (Vector Laboratories) and allowed to settle from a minimum of 2 hours to a maximum of 16 hours (overnight). Superfrost Plus slides are designed to hold tightly to cell membranes, so cells attached to these slides can be subjected to days-long labeling protocols (e.g., HCR-FISH, antibody staining, etc.) without changing location or detaching from the slide. The adherence of cells to these slides also allows for staining methods that are not simultaneously compatible to be performed sequentially with specific, individual cells being confidently reimaged throughout the sequence.

To describe their cytological profile, stem cells were identified using Piwi1 antibody staining. Antibody staining and cytological staining are not compatible when used simultaneously for two reasons: 1) cytological stains, though intended to be used colorimetrically, are broadly fluorescent and interfere with the fluorescent signal of the secondary antibody, and 2) the blocking agents required for antibody staining block the attachment of the basic stains used in cytological staining. To circumvent this, whole cell suspensions adhered to Superfrost Plus slides were first cytologically stained and imaged. Those same slides were then subjected to Piwi1 antibody labeling which detached the cytological stains during the blocking process, vastly reducing the fluorescent noise they cause, and labeled *Hydractinia*’s stem cells for imaging. By imaging the same cells following each staining method, it was possible to identify a stem cell based on its Piwi1 expression in one image and then locate that exact same cell in the second image to observe its cytological profile. A similar method was used to characterize the cytological profiles of other cell types using HCR-FISH, though in these cases HCR-FISH preceded cytological staining since the only incompatibility in these staining techniques comes from the broad fluorescence of the cytological stains.

HCR-FISH and Piwi1 antibody staining were conducted according to Song et al. (2025) and Chrysostomou et al. (2022), respectively. These protocols were modified slightly for use with the low volumes needed for slide-adhered samples; all wash steps were preceded by an additional 10-second wash of the same reagent. HCR-FISH was used to identify the following cell types using the listed genes: ectodermal epithelial cells (*Fat1* [HyS0048.57]), endodermal epithelial cells (*Astacin3* [HyS0078.51]), neuronal subtype A (*RFamide precursor* [HyS0013.338]), neuronal subtype B (HyS0049.55), zymogen gland cells (*Chit1* [HyS0041.99]), mucous gland cells (*Mucin-5AC* [HyS0004.446]), putative immune cells (HyS0016.300), cnidoblasts (*TXD12* [HyS0042.111]), and proliferating cells (*Histone H1.1* [HyS3103.5]). Each of these HCR-FISH markers has been previously validated to ensure they selectively label their associated cell type (Song et al., 2025). Cell type HCRs were performed using either a single HCR probe or, when possible, two HCR probes multiplexed using different fluorescent labels. As mentioned above, *Hydractinia*’s stem cells were labeled using Piwi1 antibody staining (Chrysostomou et al., 2022). Cnidocytes were not specifically labeled in this experiment because they can be easily identified by the presence of their large, characteristic capsule even when dissociated enzymatically. Antibody and HCR-FISH labeled cells were stained with Hoechst 33342 in 1x PBS at a concentration of 22 µM (10 µg ml^-1^) for 30 minutes to label nuclei and were mounted in 1x PBS using coverslips with clay feet prior to imaging.

Colorimetric cytological staining was performed using a combination of May-Grünwald (Eosin and Methylene Blue) and Giemsa (Eosin, Methylene Blue, and Azure B) stains (i.e., Pappenheim staining technique) which are well established for use in *Hydractinia* tissue sections (Müller, 1964, 1967; Müller et al., 2004; Pappenheim, 1908). Attached cells were gently washed with Sorensen’s phosphate buffer (SPB; pH 7.0); incubated in 1:1 SPB : May-Grünwald stock solution (Sigma-Aldrich 1.01424) for three minutes at room temperature in the dark; incubated in 9:1 SPB : Giemsa stock solution (Sigma-Aldrich 32884) for 10 minutes at room temperature in the dark; gently washed three times in SPB; and then mounted in SPB using coverslips with clay feet.

### Fluorescence and Pseudo-Colorimetric Imaging

Cells were imaged with a Leica Microsystems Stellaris confocal microscope using a 40x objective at 1.5x optical zoom through a 10x fixed lens. The scanning resolution was set at 2,048 x 2,048 pixels, the scan speed set to 400 lines per second, and the pinhole set to 3 AU for an effective optical section thickness of 5 µm. Following stage alignment, 2 – 5 large regions of interest (2-5 mm^2^ each) of properly confluent (i.e., relatively dense monolayer) cells were selected on the slide and imaged using the Stellaris’ automated tiling function and a manually calibrated focus polygon. These regions of interest were saved so that the cells within could be confidently reimaged after undergoing subsequent staining methods. When necessary, while imaging fluorescent Piwi1 antibody and HCR-FISH labeled samples, fluorescence lifetime measurements were used to exclude *Hydractinia* autofluorescence, which has a very short fluorescence lifetime (≲1 ns), using the Stellaris’ TauGating function.

Pseudo-colorimetric cytological images were captured via color transmitted light imaging (Collings, 2015) using the transmitted light photomultiplier tube (TL-PMT) in combination with the Stellaris’ adjustable white light laser (WLL). By setting the wavelength of the WLL laser to match a colorimetric stain’s maximum absorbance peak, it is possible to capture an image using the TL-PMT in which the presence of the stain is recorded as areas of low intensity (i.e., shadows) and its absence is recorded as areas of high intensity, with regions of intermediate staining showing intermediate intensity according to the relative concentration of the colorimetric stain (Fig. S2). This technique does not rely on nor measure reflective fluorescence but instead utilizes measurements of transmitted light as in standard colorimetric microscopy. To capture the pseudo-colorimetric images for the current study, three channels were used: 1) a blue channel with the WLL set to the wavelength of Giemsa stain’s minimum absorbance (405 nm) to act as an unbiased record of general refraction, 2) a green channel with the WLL set to the maximum absorbance of the Eosin component of the Giemsa stain (517 nm) to capture the cells’ Eosin profile, and 3) a red channel with the WLL set to the maximum absorbance of the Azure B component of the Giemsa stain (644 nm) to capture the cells’ Azure B and Methylene Blue components (Fig. S2). When stacked, these channels create a pseudo-colorimetric image that is functionally identical to those taken using standard colorimetric microscopy (Fig.1). Comparing the methods revealed that the informative aspects of the images (i.e., the location, intensity, and staining pattern of each component stain) are indistinguishable between them, with the only notable difference being the visual color of each staining component (Eosin is light brown in standard colorimetric and red-purple in pseudo-colorimetric; Azure B is deep indigo in standard colorimetric and green-blue in pseudo-colorimetric) (Fig. 1). Though the Methylene Blue component of the Giemsa stain was not imaged on its own specific channel, its visual color, maximum absorbance peak, and staining pattern are nearly identical to that of Giemsa’s other basic dye, Azure B, and its role in the Giemsa staining solution is to simply deepen the color of the acidic structure labeled by Azure B. In other words, the staining profile of Methylene Blue was captured on the red channel along with Azure B, with which it is largely redundant.

**Figure 1.**
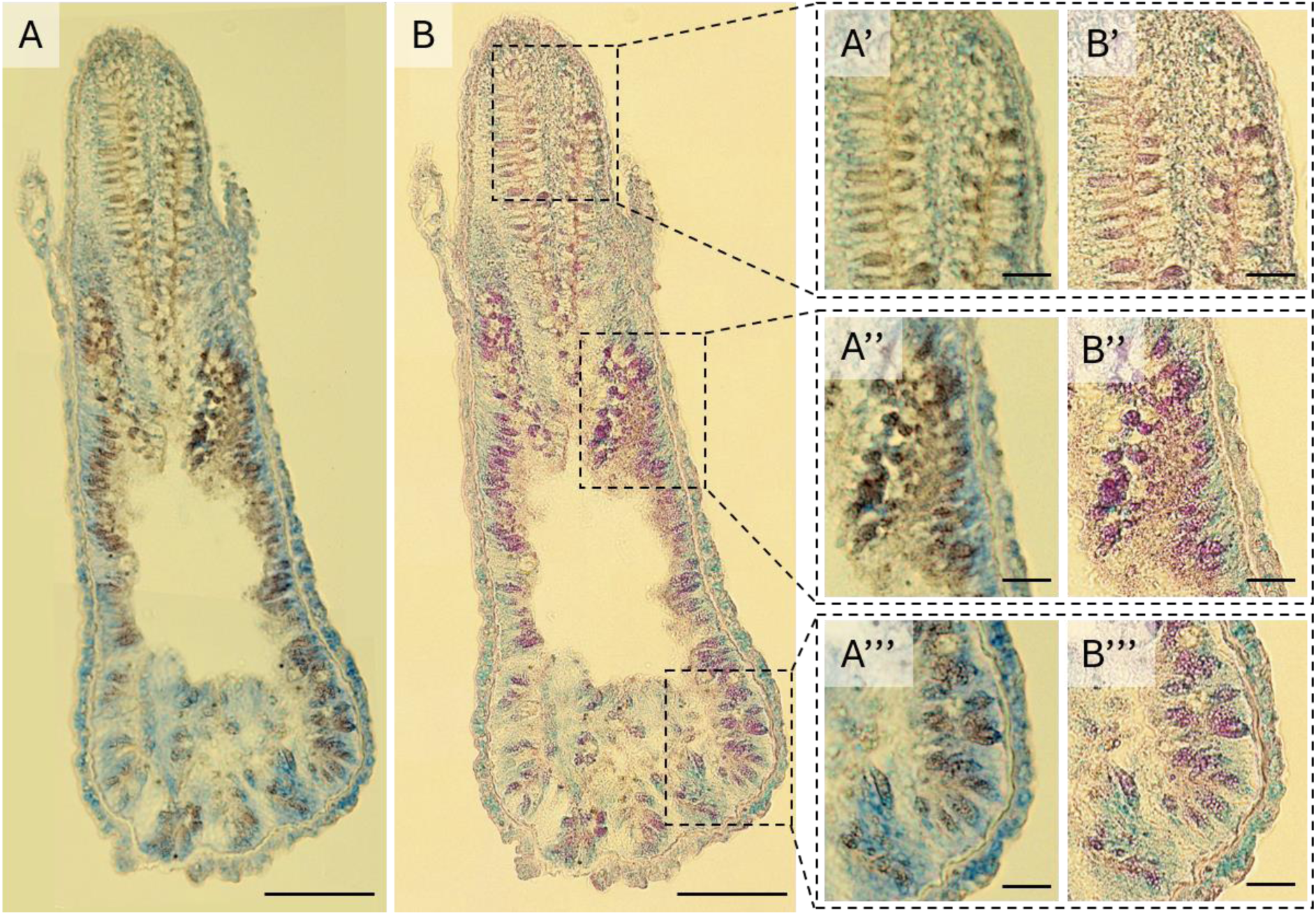
Comparison of (A) colorimetric and (B) pseudo-colorimetric images of a single *Hydractinia* feeding polyp tissue section stained using the May-Grünwald/Giemsa staining technique. Large insert scale bars are 100 µm and small insert scale bars are 20 µm.

### Mitomycin C Treatment

To selectively deplete proliferative stem cells, *Hydractinia* colonies were exposed to the cytostatic, DNA-alkylating agent Mitomycin C (MitC; Santa Cruz Biotechnology) (Müller, 1967, 1968; Müller et al., 2004). Prior to treatment, colonies were not fed for three days. For each colony treated, the microscope slide on which that colony resided was first split in half via etching and careful breaking, with one half being randomly assigned to the mitomycin treatment condition and the other half being assigned as a negative control. The two halves were then placed into 60 mm glass petri dishes and submerged in 9.4 ml of artificial seawater at 30 ppt. Then, 600 μl of 1 mM MitC solution in Milli-Q H_2_O was added to the dish containing to the treatment colony to achieve a final incubation volume and concentration of 10 ml of 60 μM MitC. To account for the slight difference in salinity brought on by the addition of the MitC solution, 600 ul of Milli-Q water was added to the dish containing the negative control colony to achieve a final incubation volume of 10 ml. Both colonies were then incubated at 20°C in the dark for 18 hours. Following incubation, each colony was separately rinsed in artificial seawater and placed into a holding tank under the conditions outlined in the Animal Husbandry section above for three days before cell dissociation and flow cytometry analysis (Fig. S3).

### Preparation of Single-Cell Suspension for FACS Analysis

Cellular suspensions were characterized using the following fluorescent dyes: DAPI (Sigma-Aldrich; excitation 358 nm; emission 461 nm) was used at a final concentration of 1.43 μM (0.25 μg ml^-1^) to exclude dead cells; DRAQ5 (Miltenyi Biotec; excitation 601 nm; emission 699 nm) was used at a final concentration of 15 µM to stain nuclear DNA; Pyronin Y (Thermo Scientific; excitation 548 nm; emission 566 nm) was used at a final concentration of 16.5 µM (5 µg ml^-1^) to stain cellular RNA; and Calcein AM (Invitrogen; excitation 495 nm; emission 515 nm) was used at a final concentration of 3.2 µM (2 µg ml^-1^) to determine the cellular volume of viable cells. Cell suspensions were incubated with these stains for 45 minutes at room temperature in the dark. Following incubation, cells were pelleted at 800 rcf for 5 minutes at 4°C and then resuspended in ice-cold CMFASW (Fig. S1). Preliminary control experiments demonstrated that live *Hydractinia* cells efflux DRAQ5 and Pyronin Y quickly enough to alter cytometric profiles taken over the course several hours (e.g., during lengthy sorting experiments), so following resuspension in ice-cold CMFASW both of these stains were added back into solution to a final concentration of 10 µM each. This concentration, while not high enough to achieve stoichiometric binding during initial staining, is adequate to counteract stain efflux throughout analysis and low enough to avoid analytical artifacts that sometimes occur when DRAQ5 and Pyronin Y are combined at high concentrations. Following staining, cell suspensions were kept on ice in the dark until flow cytometry analysis.

### Flow Cytometry Analysis and Cell Sorting

Flow cytometry analysis and cell sorting were performed using a three-laser (405 nm violet, 488 nm blue, and 561 nm yellow-green) BD FACSMelody cell sorter. DAPI fluorescence intensity was measured on the violet laser’s 488/45 detector, Calcein on the blue laser’s 527/32 detector, and Pyronin Y and DRAQ5 on the yellow-green laser’s 528/15 and 697/58 detectors, respectively. Prior to analysis, single-stain compensation controls, double-stain controls, and fluorescence-minus-one (FMO) controls were collected, along with a cnidarian-specific control comprised of isolated cnidocyte capsules and nuclei to ensure that the chosen stains did not interact with cnidocyte capsules in unexpected ways. Compensation was required to account for the minimal spillover of Pyronin Y into the DRAQ5 detector. DAPI, Calcein, and Pyronin Y were bi-exponentially voltrated; side scatter (SSC) was logarithmically voltrated; and DRAQ5 and forward scatter (FSC) were linearly voltrated. Dead cells were excluded using DAPI area x Calcein area and doublet exclusion was performed using DRAQ5 area x DRAQ5 width (Fig. 2 A and B). Cnidocytes are identifiable based on their DNA content (2n) and high SSC, and were therefore able to be excluded using DRAQ5 area x SSC area (Fig. 2 C).

**Figure 2.**
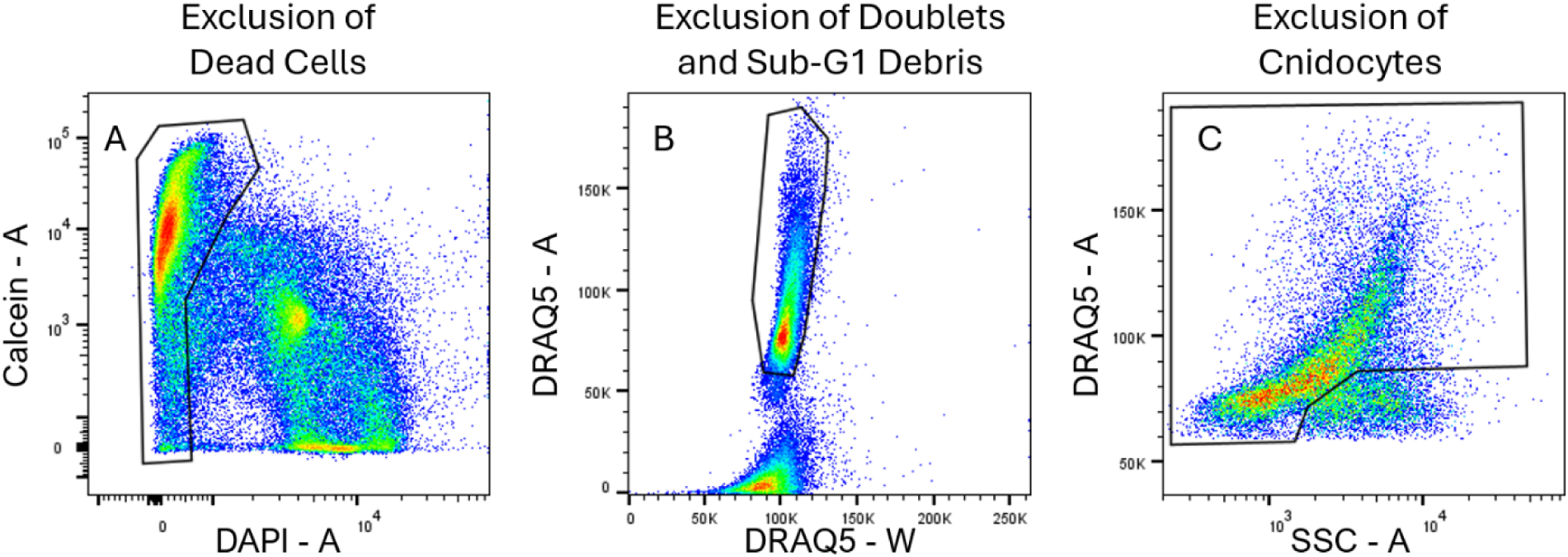
Basic gating strategies to (A) exclude DAPI^+^ dead cells, (B) exclude doublets with high DRAQ5 width-to-area ratios and cellular debris with sub-2n DRAQ5–A intensities, and (C) exclude cnidocytes based on their 2n DNA content, as determined by DRAQ5–A intensities, and high SSC–A values.

Following the three basic gates outlined above, stem cells were identified via comparisons of MitC treated and untreated control samples. MitC selectively depletes stem cells in *Hydractinia* (Müller, 1967, 1968; Müller et al., 2004), so by gating the cell population that was depleted in the MitC sample but not depleted in the control sample (i.e., subtractive gating), it was possible to create a nested series of two gates that effectively isolated the depleted population of interest (called the primary and secondary ‘stem gates’ from here on; see Results for more detail) (Fig. 3). The cells within the stem gates were then sorted from the untreated control sample. During sorting, temperature control was used to keep both the sample and the sorted cells at 4°C to minimize cell death. A 100 µm sort nozzle and BD FACSFlow sheath fluid were used, and cells were sorted into 5 ml polypropylene test tubes precoated with 2% BSA in CMFASW containing 2 ml of CMFASW catch buffer. The sorted cells were then sequentially stained and imaged using May Grünwald/Giemsa staining and Piwi1 antibody staining as outlined above to determine identity (Fig S3).

**Figure 3.**
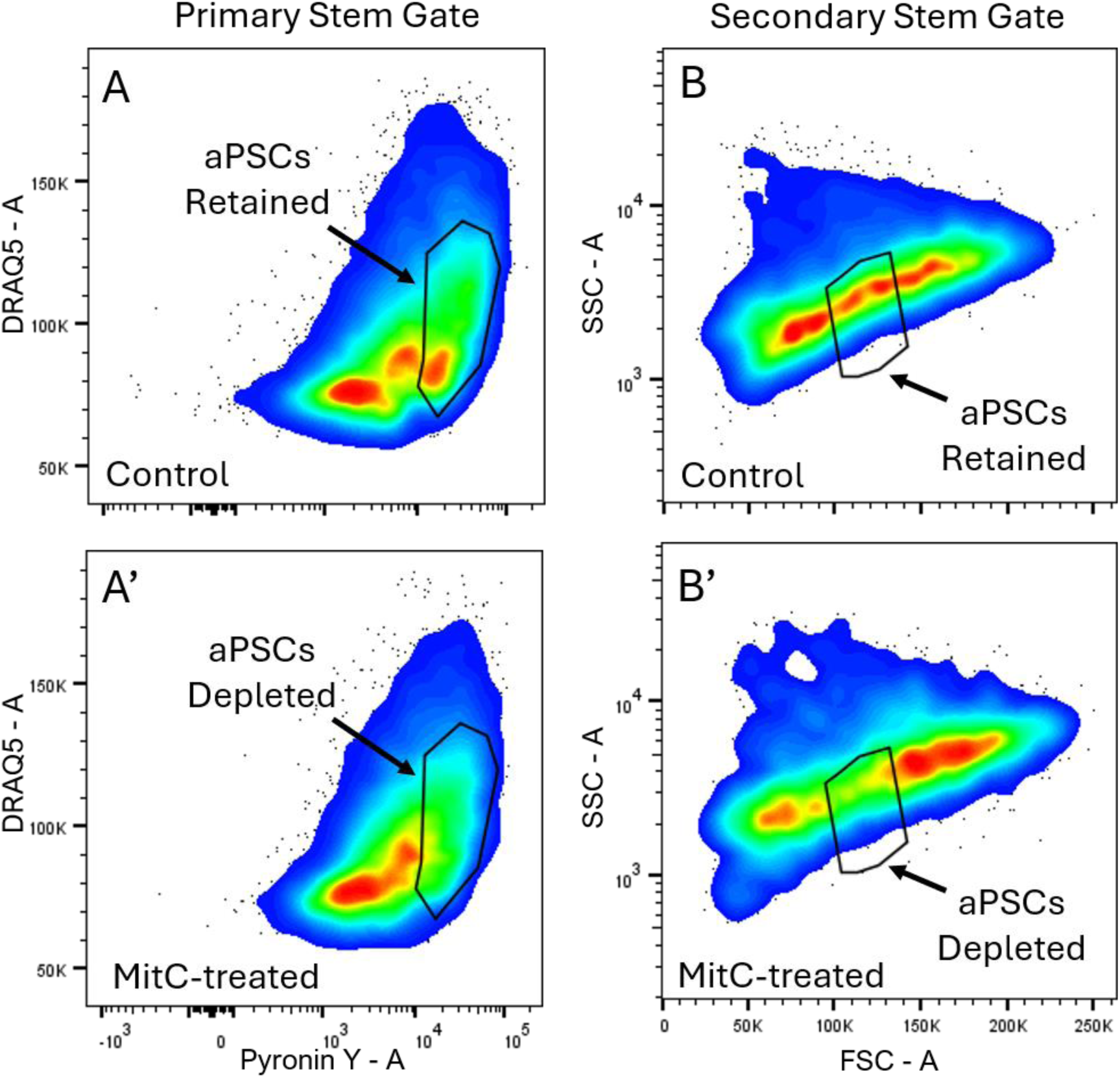
Flow cytometry plots showing the (A) primary stem gate and (B) secondary stem gate in both control (no apostrophe) and drug-treated (apostrophe) conditions. Heat maps indicate high event concentrations in red to low event concentrations in blue. Outlying events are indicated as dots.

### Statistical Analysis of Stem Gates

To assess the effectiveness and consistency of the MitC treatment and the reproducibility of the primary and secondary stem gates, data were collected from seven MitC treated colonies and five untreated colonies over the course of three sampling periods. Due to the drift in fluorescence intensity readings that naturally occurs over time in flow cytometers, all detectors were re-voltrated and all gates were redrawn for each sampling period, with the same settings and gates being used for each sample recorded within a single sampling period. For each sample, the percentage of total live cells within the primary and secondary stem gates (percentage of cells on-target for each gate) were recorded as was the percentage of total live cells outside each stem gate (percentage of cells off-target for each gate).

The following statistical analyses were conducted in R v4.5.3 (R Core Team, 2026) using RStudio (Posit Team, 2025). To assess if the cells within the primary and secondary stem gates were significantly depleted under MitC conditions compared to control conditions, a linear mixed effects model (model 1) with logit-transformed percentage of total cells within each gate as the dependent variable; treatment group (MitC/control) and gate identity (primary/secondary) as fixed effects; and sample (to account for repeated measures) nested within sampling period (to account for differences in instrument settings) as random effects was created and applied (lme4 package; Bates et al., 2015). To assess whether or not cells within the stem gates (on-target) were statistically depleted compared to cells outside of the stem gates (off-target) under MitC conditions compared to control conditions, a second linear mixed effects model (model 2) with logit-transformed percentage of total cells as the dependent variable; treatment group (MitC/control), gate identity (primary/secondary), and gate status (on-target/off-target) as fixed effects; and sample nested within sampling period as random effects was created and applied (lme4 package; Bates et al., 2015). Assumptions of residual normality and homoscedasticity were tested and met for both models. Significance of the models’ main effects and interaction terms were determined using Type III ANOVA (lmerTest package; Kuznetsova et al., 2017), and post-hoc pairwise comparisons were tested using a false discovery rate p-value adjustment (grafify package; Shenoy, 2021).

### Preparation of formalin-fixed paraffin-embedded tissue sections

Formalin-fixed paraffin-embedded tissue sections were used to assess the comparability of standard colorimetric and pseudo-colorimetric imaging techniques. *Hydractinia* colonies were relaxed in 700 mM MgCl (40 g l^-1^) in 15 ppt filtered artificial seawater. Feeding polyps were dissected and fixed in 4% paraformaldehyde in 30 ppt filtered artificial sweater for 2 hours on a rocker at room temperature. Feeding polyps were then dehydrated via serial 10 min baths in an increasing ethanol series (50 – 100%) and incubated twice in 100% xylene for 10 min each. Polyps were then carefully embedded and positioned in liquid paraffin wax (melt point 56-58°C; Epredia B1002490) in polyethylene molds (Ted Pella Inc 27110) placed on a heating plate to ensure that the wax did not solidify during embedding. After allowing polyps to incubate in the liquid wax for 10 min on the heating plate, the molds were removed from heat, the wax was allowed to reach room temperature, and then the molds were peeled away. The back side of the wax blocks were then quickly melted with a broad knife and flame to be attached to a microtome cassette. The blocks were then stored on ice to further solidify the wax before they were loaded into a Leica Microsystems RM 2235 microtome and sectioned at a 5 µm thickness. Sections were mounted on Superfrost Plus slides via a water bath (i.e., sections were floated on the water surface, positioned to the edge of a vertically submerged slide, and mounted as the slide was vertically removed from bath), and allowed to fully dry overnight. Slides were deparaffinized and rehydrated via a series of 5 min baths in a decreasing xylene/ethanol series (100% xylene – pure diH_2_O) and were allowed to completely dry before staining. A separate, detailed protocol of this technique is available (Lane, 2026).

## RESULTS

### Cytological Characterization of Cell Types

From our analysis, we show that it is possible to discriminate between live-dissociated *Hydractinia* cell types through cytological staining and cell diameter measurements. Representative images and average diameters for each cell type are presented in Figure 3 and general descriptions are outlined below. Table S1 contains short-hand descriptions, average sizes, and diagrams of simplified cytological profiles for each cell type and is intended as a quick reference for future assay execution. Significant differences in diameter between cell types were determined via a single One-way ANOVA and post-hoc Tukey’s HSD (Fig. S3).

Stem cells, identified via Piwi1 positivity and *Histone H1.1* positivity (Fig. 4A & B), as well as their immediate cnidoblast progenitors (*TXD12*; Fig. 4C), share a similar and distinct cytological profile. These undifferentiated cell types are dark in color and have a high nuclear-to-cytoplasmic ratio compared to other cell types. Their cytoplasmic and nuclear compartments are clearly delineated, with the cytoplasm appearing dark blue and their nucleus taking a redder purple hue. These cell types do not differ significantly in size from one another and are relatively small compared to other cell types (Piwi1^+^: 5.92 ± 0.48 µm; *Histone H1.1*^+^: 6.15 ± 0.59 µm; *TXD12*: 5.73 ± 0.71 µm; n = 22 for each).

**Figure 4.**
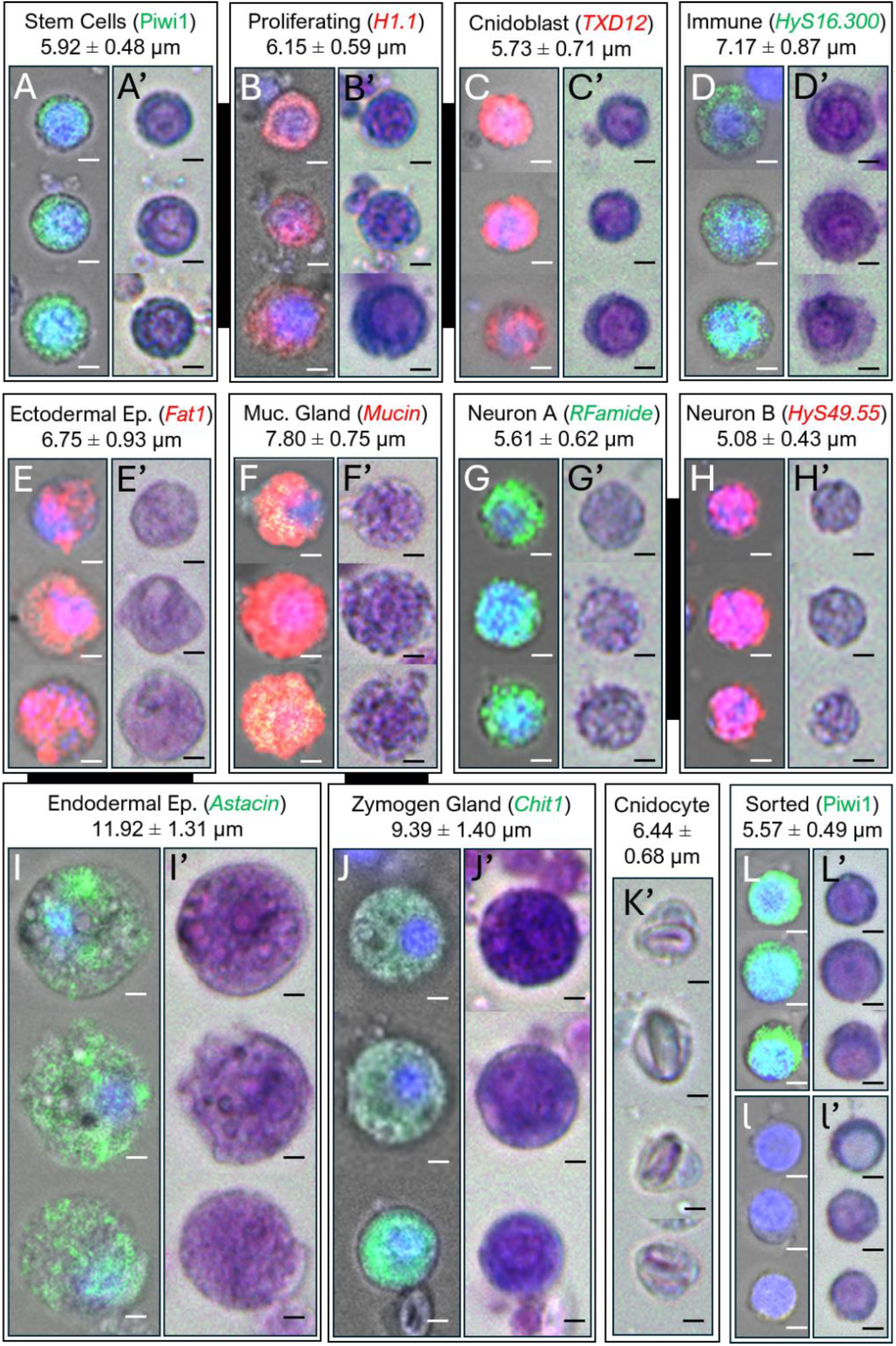
Enzymatically dissociated (A) stem cells, (B) proliferating cells, (C) cnidoblasts, (D) putative immune cells, (E) ectodermal epithelial cells, (F) mucous gland cells, (G) neuronal subtype A, (H) neuronal subtype B, (I), endodermal epithelial cells, (J) zymogen gland cells, (K) cnidocytes, and both (L) Piwi1^+^ and (l) Piwi1^-^ sorted cells stained with expression-based makers (no apostrophe) and cytological stains (apostrophe). Similar cell types (e.g., gland cells) are linked by black connections. Cell type and marker identity labels are listed with mean cellular diameter ± standard deviation above images. All cells scaled identically and scale bars are 2 µm. *HyS001c.300* and *HyS004S.55* are abbreviated.

Immune cells (*HyS001c.300*) have a clearly delineated nuclear envelope, similar to the undifferentiated cells above (Fig. 4D). However, immune cells can be confidently discriminated from these undifferentiated cell types by a reddish-purple coloration throughout all cellular compartments, their relatively smaller nuclear-to-cytoplasmic ratio, their variable granularity, and their significantly larger size (7.17 ± 0.89 µm, n = 22) (One way ANOVA, F = 103.9, post-hoc Tukey HSD, p < 0.05).

Ectodermal (*Fat1*) and endodermal (*Astacin*) epithelial cells were characterized by Eosin heavy staining, taking on a lighter, redder staining profile than other cell types of similar size and lacking a clearly stained nucleus (Fig 4E & I). Endodermal epithelial cells were significantly larger (11.92 ± 1.39 µm, n = 10) than any other cell type (One way ANOVA, F = 103.9, post-hoc Tukey HSD, p < 0.05) and often had vacuoles of various sizes throughout their cytoplasm. Ectodermal epithelial cells we significantly larger (6.75 ± 0.95 µm, n = 22) than neurons and smaller than gland cells and endodermal epithelial cells (One-way ANOVA, F = 103.9, post-hoc Tukey HSD, p < 0.05). Ectodermal epithelial cells have a similar size to immune cells but lack the clearly visible nuclear envelope of immune cells, generally take on a lighter staining profile, and are slightly smaller on average.

Gland cells are some of the largest cells with zymogen gland cells (*Chit1*) being significantly larger (9.39 ± 1.44 µm, n = 22) than all but endodermal epithelial cells, and mucous gland cells (*Mucin*) being significantly smaller (7.80 ± 0.77 µm; n = 22) than zymogen gland cells but larger than all other cell types besides immune cells (One-way ANOVA, F = 103.9, post-hoc Tukey HSD, p < 0.05) (Fig. 4F & J). Mucous gland cells are recognizable by their relatively light staining profile and numerous granules, while zymogen gland cells can be identified by their large size and dark purple staining profile. Both gland cell types typically lack a clearly visible nucleus, but nuclei are occasionally visible with a low nuclear-to-cytoplasmic ratio.

Both neuronal subtypes A (*RFamide*) and B (*HyS004S.55*) are small and sparsely stained with some granules and no strong nuclear staining (Fig. 4G & H). Neurons were the smallest cells on average with subtype B (5.08 ± 0.44 µm, n = 22) being smaller on average than subtype A (5.61 ± 0.64 µm, n = 22). Though these cell types did not differ significantly in size from one another or from the stem cell group described above, they were significantly smaller than any other differentiated cell type (One way ANOVA, F = 103.9, post-hoc Tukey HSD, p < 0.05).

Finally, as mentioned previously, cnidocytes can be identified by the presence of their conspicuous, refractory capsule (Fig. 4K) even in the absence of any cytological stain. These cells are of an intermediate size (6.44 ± 0.70 µm).

### Flow Cytometry Analysis and Characteristics of Isolated Cell Population

Following the creation of the three basic flow cytometry gates outlined above (dead cell exclusion, doublet/debris exclusion, cnidocyte exclusion) (Fig. 2), the MitC-depleted stem cell population was isolated through the use of a primary and secondary stem gate (Fig. 3). The primary stem gate utilized the DRAQ5-A x Pyronin Y-A axes and encompassed a population of RNA-high cells depleted by MitC exposure that extended through the DRAQ5-A axis from 2n – 4n DNA content (Fig 3A & A’). The secondary stem gate utilized the FSC-A x SSC-A axes to further hone this population of RNA-high cells to isolate a population of low SSC, intermediately sized cells that were selectively depleted in MitC-treated samples (Fig3B & B’). The populations within both gates were significantly depleted in MitC-treated samples compared to control samples (Model 1, Type III ANOVA, treatment x gate identity interaction, F = 8.74, p = 0.014, post-hoc pairwise comparisons p < 0.05), with the primary gate being depleted by ∼25% in MitC-treated samples (control: 37.00 ± 10.99% of total cells within gate; MitC-treated: 28.42 ± 8.49% of total cells within gate) and the secondary gate being depleted by ∼55% in MitC-treated samples (control: 7.65 ± 3.44% of total cells within gate; MitC-treated: 3.44 ± 1.58% of total cells within gate) (Fig. 5). Further, the cell population within these gates was shown to be selectively depleted in MitC-treated samples compared to cell populations outside the gates which were not depleted, and this difference was significant according to statistical analysis (Model 2, Type III ANOVA, treatment x gate status interaction, F = 6.78, p = 0.013, post-hoc pairwise comparisons p < 0.05) indicating that the final gates were precisely shaped to include the majority of the MitC-depleted population (Fig. 6). The DRAQ5 fluorescence intensity of the final gated population was used to estimate the cell cycle status of the cells and indicated that 82.66 ± 3.84% of the final population was in the G1 stage and 17.34 ±3.84% was in the S/G2 stage. Cell cycle dynamics did not differ between control and MitC-treated samples (Students T-test of logit-transformed G1 percentages, t = 1.84, p = 0.097).

**Figure 5.**
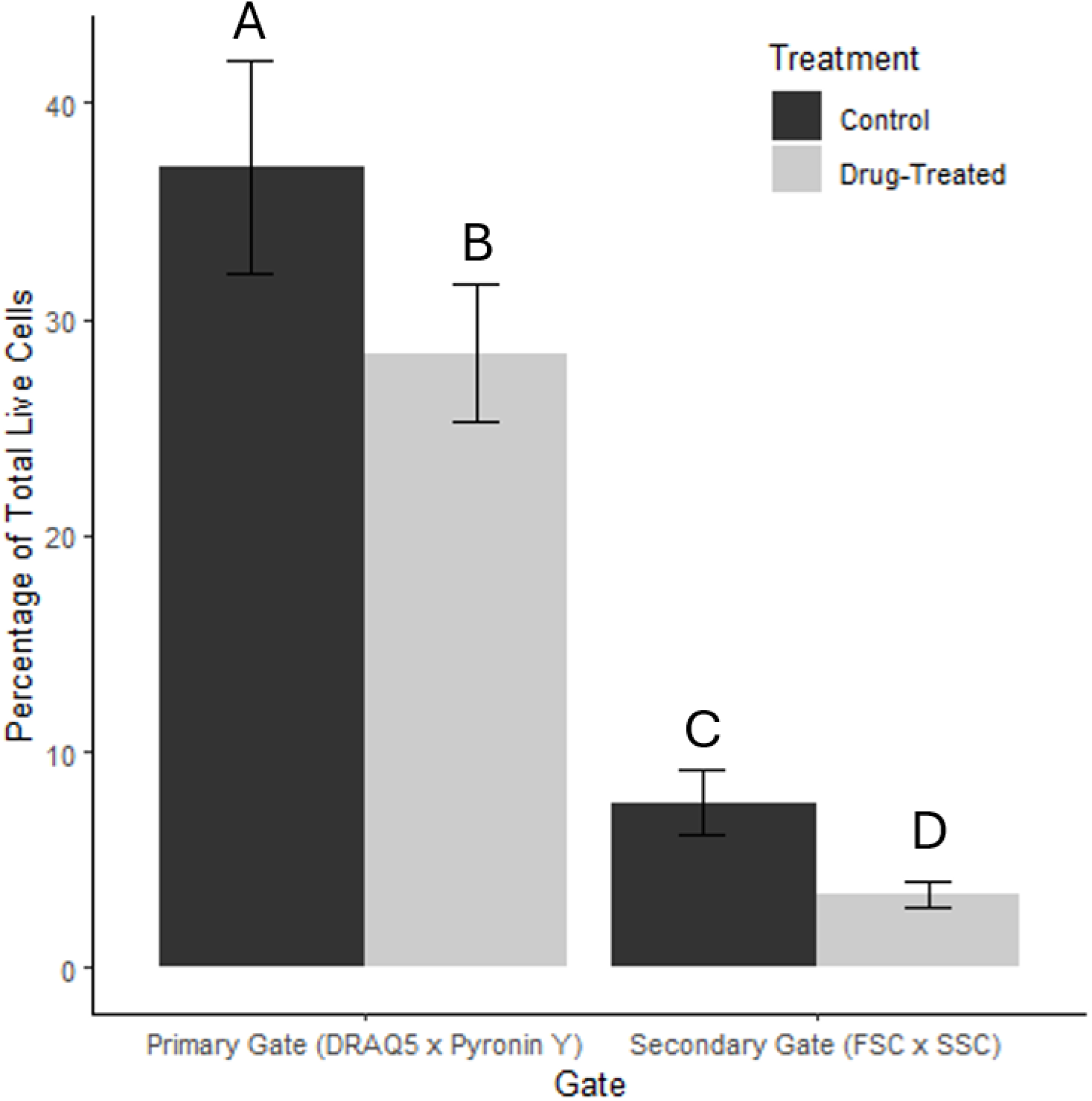
Plot showing differences in the percentage of total cells within the primary and secondary stem gates between control (dark grey) and drug-treated (light grey) conditions. Significant differences according to Model 1 type III ANOVA and post-hoc pairwise analysis with α = 0.05 are indicated in standard compact letter display format. Error bars show standard error.

**Figure 6.**
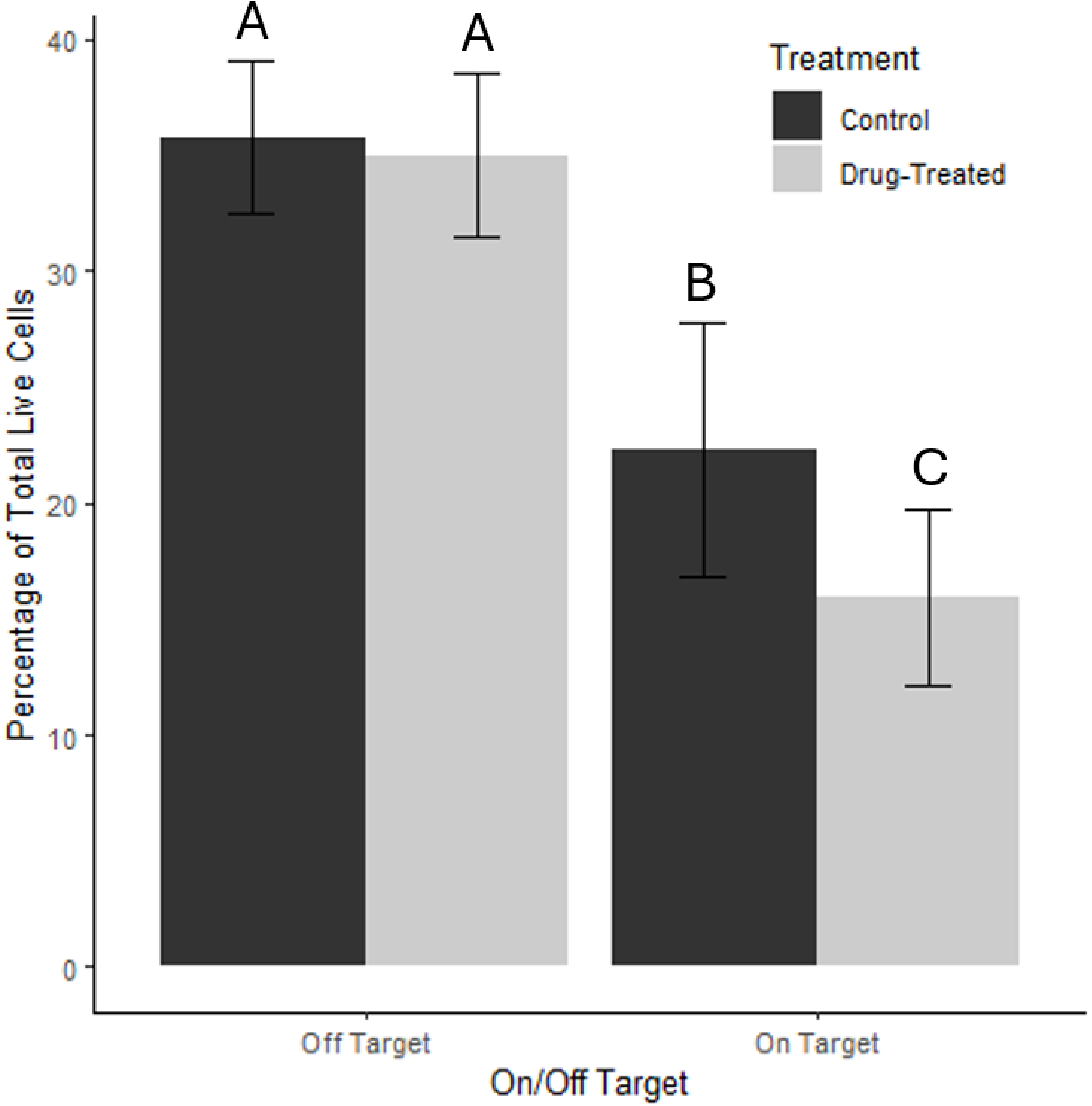
Plot showing differences in the percentage of total cells within stem gates (on-target) and outside of stem gates (off-target) between control (dark grey) and drug-treated (light grey) conditions. Significant differences according to Model 2 type III ANOVA and post-hoc pairwise analysis with α = 0.05 are indicated in standard compact letter display format. Error bars show standard error.

The isolated cell population was comprised mostly (>80%) of cells that displayed the cytological characteristics of stem cells and cnidoblasts outlined above, showing clearly delineated cytoplasmic and nuclear compartments with the cytoplasm appearing dark blue and their nucleus taking a redder purple hue (Fig. 4L & l). The isolated population did not differ significantly in size (5.83 ± 0.52 µm, n = 100) from stem cells (Piwi1 and *Histone H1.1*) or cnidoblasts (*TXD12*) described above but was significantly smaller than all differentiated cell types besides neurons (One way ANOVA, F = 103.9, post-hoc Tukey HSD, p < 0.05) (Fig. S3). Of the <20% of the isolated cell population that did not show clearly identifiable cytological characteristics of undifferentiated cells, ∼15% were apparently destroyed and could not be identified, and the remaining ∼5% appeared to be developing cnidocytes with small capsules or small ectodermal epithelial cells. Roughly 15% of the intact isolated population was Piwi1^+^ according to antibody labeling, which was a ∼10x increase in the density of such cells compared to whole cell suspension in this study.

## DISCUSSION

### Cytological assay

The cytological assay developed for this project allowed for the simultaneous assignment of functional identity to all major cell types in live suspension (Fig. 4 and Table S1). This single assay is sufficient to discriminate between zymogen and mucous gland cells, endo-and ectodermal epithelial cells, immune cells, cnidocytes, neurons, and undifferentiated cells, though it is limited by its inability to differentiate type A from type B neurons and aPSCs from their immediate, still-differentiating progenitors. Though validated marker gene expression-based labels exist for all major cell types in *Hydractinia*, this cytological assay may be more appropriate for certain use-cases for the following reasons: (1) the assay is extremely efficient compared to cell-specific labeling, being much faster (i.e., taking 2-3 hours rather than the 2-3 days required for antibody labeling or HCR-FISH), more methodologically simple, cheaper, and it utilizes generically available staining solutions, (2) the assay has the ability to simultaneously identify all major cell types without *a priori* knowledge of cell solution composition and without complex fluorescence multiplexing, and (3) the assay may mitigate issues of partial/extra-population labeling that can occur when marker expression varies within/among cell types. With this assay is possible to subsample a live *Hydractinia* cell suspension and discern the cell types present in a short enough time frame to proceed to downstream applications with the remainder of the live cell suspension. Our findings concerning the cytological profiles of live dissociated cells are largely in agreement with the histological profiles previously described for these cells in whole mount and tissue section analyses (Müller, 1964, 1967; Müller et al., 2004).

### Sorted cell population

The following lines of evidence support the claim that the sorted population is comprised of *Hydractinia* aPSCs: (1) the sorted population was selectively depleted by the administration of Mitomycin C, a DNA-alkylating agent known to selectively deplete *Hydractinia* aPSCs (Müller, 1967, 1968; Müller et al., 2004); (2) the sorted population had a high RNA concentration based on Pyronin Y fluorescence intensity and were proliferative based on DRAQ5 cell cycle estimation, both of which are general characteristics of stem cells (Molinaro et al., 2021; Rhee and Bao, 2009; Rumman et al., 2015; Zakrzewski et al., 2019); (3) the large majority of cells in the sorted population had a size and cytological profile indicative of *Hydractinia* aPSCs according to the cytological assay developed in this study; and (4) cells with high levels of Piwi1 expression, which is a known aPSC marker in *Hydractinia*, were ∼10x enriched in the sorted population compared to whole cell suspension, comprising roughly 15% and 1.5%, respectively. The percentage of total cells within the secondary stem gate of untreated control samples (7.65 ± 3.44%) is higher than previous estimates of total aPSC percentages for *Hydractinia* (Chrysostomou et al., 2022; DuBuc et al., 2020), though these previous estimates were based entirely on Piwi1^+^ aPSCs and did not account for variable expression of Piwi1 in *Hydractinia* aPSCs (Song et al., 2025; Waletich et al., 2024). The percentage of cells that fell into the secondary stem gate and were also Piwi1^+^ (∼1.1% of total cells according to post-sort antibody staining) is nearly identical to previous estimates of total Piwi1^+^ aPSC percentages (Chrysostomou et al., 2022; DuBuc et al., 2020). This would indicate that the vast majority of the Piwi1^+^ aPSC population was included in our final sorting gate. As previously stated, our cytological assay cannot differentiate between aPSCs and their immediate, differentiating progenitors. Given the lack of cytological differences and the fact that progenitors are likely to be depleted in MitC-treated colonies as well as aPSCs, it is not currently possible for us to reject the possibility that our sorted population is comprised of an unknown ratio of aPSCs and their immediate progenitors, rather than a pure population of aPSCs. For some downstream applications, it may be advantageous to isolate a more inclusive population of stem cells that includes immediate progenitors. Further functional experimentation (e.g., sorted cell transplantation) and transcriptional analyses (e.g., single-cell RNA sequencing) are warranted to fully characterize the identity and stemness of the isolated population to determine if a modified approach would yield a pure population of aPSCs.

Our cell cycle estimates for the sorted aPSC population (G1: 82.66 ± 3.84%; S/G2: 17.34 ±3.84%) differs substantially from a previous study by Chrysostomou et al. (2022) which indicated that nearly 100% of *Hydractinia* aPSCs fall within S/G2 phase. The reason for this discrepancy cannot be definitively stated with current data, but the difference in cell cycle estimates may be due to Chrysostomou et al. (2022) utilizing a Piwi1-associated transgenic line to identify *Hydractinia* aPSCs rather than an expression-agnostic approach like the one used in the current study. One hypothesis is that Piwi1 is expressed mostly in aPSCs in the S/G2 phase. This hypothesis fits our understanding that Piwi1 is expressed only in a subset of *Hydractinia* aPSCs (Song et al., 2025; Waletich et al., 2024), and correlates with the findings of the current study where a similar number of aPSCs show Piwi1 positivity (∼15%) as are found in the S/G2 phase of the cell cycle (17.34 ±3.84%). Alternatively, the high number of G1 phase cells found in the sorted population in the current study could be explained by the inclusion of immediate aPSC progenitors in our stem gates. Additional research on this topic is needed to conclusively address this discrepancy in findings.

### Note on the Pseudo-Colorimetric Imaging Technique

While the color transmitted light imaging technique (Collings, 2015) used to create the pseudo-colorimetric images in the current study generates images that are functionally identical to standard colorimetric images (i.e., both contain the same type and quality of information and can be used for the same purposes) (Fig. 1 and Fig. S2), its usage is not widespread and, to our knowledge, has not been previously used in conjunction with the fine-scale laser wavelength adjustment made possible by modern WLLs to target specific colorimetric stains. The benefits of this technique include its usefulness in creating precise colorimetric images using microscopy equipment that is otherwise incapable of standard colorimetric imaging (e.g., in confocal microscopy), its ability to measure specific colorimetric stain intensity when laser wavelengths are matched to that stain’s absorbance peak, and its ability to separate concurrent colorimetric stain signals into separate channels rather than having all concurrent colorimetric signals mixed within a single standard colorimetric image. In addition, though not utilized in this study, it is possible to invert the values of absorbance peak-matched channels to create images in which regions of stain concentration appear as bright spots (similar to how fluorescent signal is typically displayed), rather than as shadows, which may be preferrable in certain use-cases (Fig S2). As many institutions invest in cutting edge fluorescence microscopy equipment (such as the Leica Microsystems Stellaris confocal microscope used in the current study), this technique may be useful for researchers who wish to utilize modern instrument precision and ease-of-use functions, perform side-by-side colorimetric and fluorescence imaging, or otherwise lack access to standard colorimetric imaging options.

Given the uncommonness of the technique, it is important to highlight the fact that the apparent colors of the pseudo-colorimetric images are arbitrary (Fig. S4). Each of the three channels that comprise the pseudo-colorimetric images in this study is a single-band 2D raster containing transmitted light intensity values ranging from 0 – 256 recorded by the TL-PMT for every pixel, as standard in 24-bit digital microscopy (Wallace et al., 2018). These values are then assigned an arbitrary color based on the look-up table (LUT) selected by the user. In the current study, general refraction intensities were displayed with a blue LUT, Eosin absorbance with a green LUT, and Azure B/Methylene Blue absorbance with a red LUT. These LUTs were chosen because blue, green, and red combined to make white in areas of low absorbance (i.e., these LUTs created a white background), and they were assigned as they were to the specific channels because this yielded the pseudo-colorimetric image that was the most visually similar to the standard colorimetric profile of the Giemsa stain out of all the potential LUT assignment permutations (Fig. S4). This sort of arbitrary LUT color selection is common in fluorescence microscopy which frequently utilizes reflected light detectors or high-sensitivity black and white cameras that create single-band 2D rasters of fluorescence intensity (Wallace et al., 2018). In either case, researchers that frequently perform fluorescence microscopy will likely already be familiar with the assignment of LUTs to increase the contrast and legibility of their images.

With this in mind, we offer the following advice for researchers hoping to utilize our cytological assay to assign cell types in live-dissociated *Hydractinia* cell suspension: (1) if creating your own pseudo-colorimetric images to perform the assay, use the same channels, laser wavelengths, and LUTs as those presented in this manuscript to ensure that your final images are easily comparable with our own, and (2) if capturing standard colorimetric images of Giemsa cytological staining in *Hydractinia*, expect the reds and blues of our images to appear more translucent brown and darker indigo, respectively.

## Conclusions

Here we have described an expression-agnostic FACS technique for the isolation of a live *Hydractinia* cell suspension highly enriched with aPSCs, demonstrated its specificity, and validated, to the best of our abilities, the identity of the sorted cells through cytology and antibody staining. We identified the aPSC population by leveraging known biological aspects of *Hydractinia* aPSCs, namely their high RNA content and sensitivity to the DNA-alkylating agent Mitomycin C. Though MitC was used to initially identify the aPSC population, experimental administration of this drug is not necessary to isolate the described population for future use. Additionally, though Calcein AM was included in our FACS staining panel, its role in our basic gates (Fig. 2) is not strictly necessary and it could be removed or replaced with a different fluorophore with similar excitation/emission. Ultimately, the FACS technique we have described will enable experimental cell transplantation, cell culture, and spheroid development using isolated *Hydractinia* aPSCs, and will further our understanding of the maintenance and control of these cells during homeostasis, regeneration, wound healing, and aging in *Hydractinia.* Additionally, we have described a cytological assay which enables the simultaneous identification of all major cell types in live cell suspension for *Hydractinia.* This assay was useful in the validation of our FACS technique for sorting the aPSC population and has the potential to expedite future studies requiring cell type identification in *Hydractinia* and enable studies that are logistically impossible under the time frames required for expression-based labeling.

## ACKNOWLEDGEMENTS

Thanks to Dr. Danielle de Jong for assistance in developing the antibody and HCR-FISH staining protocols for dissociated cells, to Dr. Jessica Farrell for sharing her histological experience during assay development, and to Dr. James Strother for discussions concerning the color transmitted light imaging technique.

## AUTHOR CONTRIBUTIONS

Z.L. and C.S. conceived and designed the study; wrote the main manuscript text or substantively revised it; and contributed to the methodology, acquisition of data and analysis, and interpretation of data.

---

**Figure S1.**
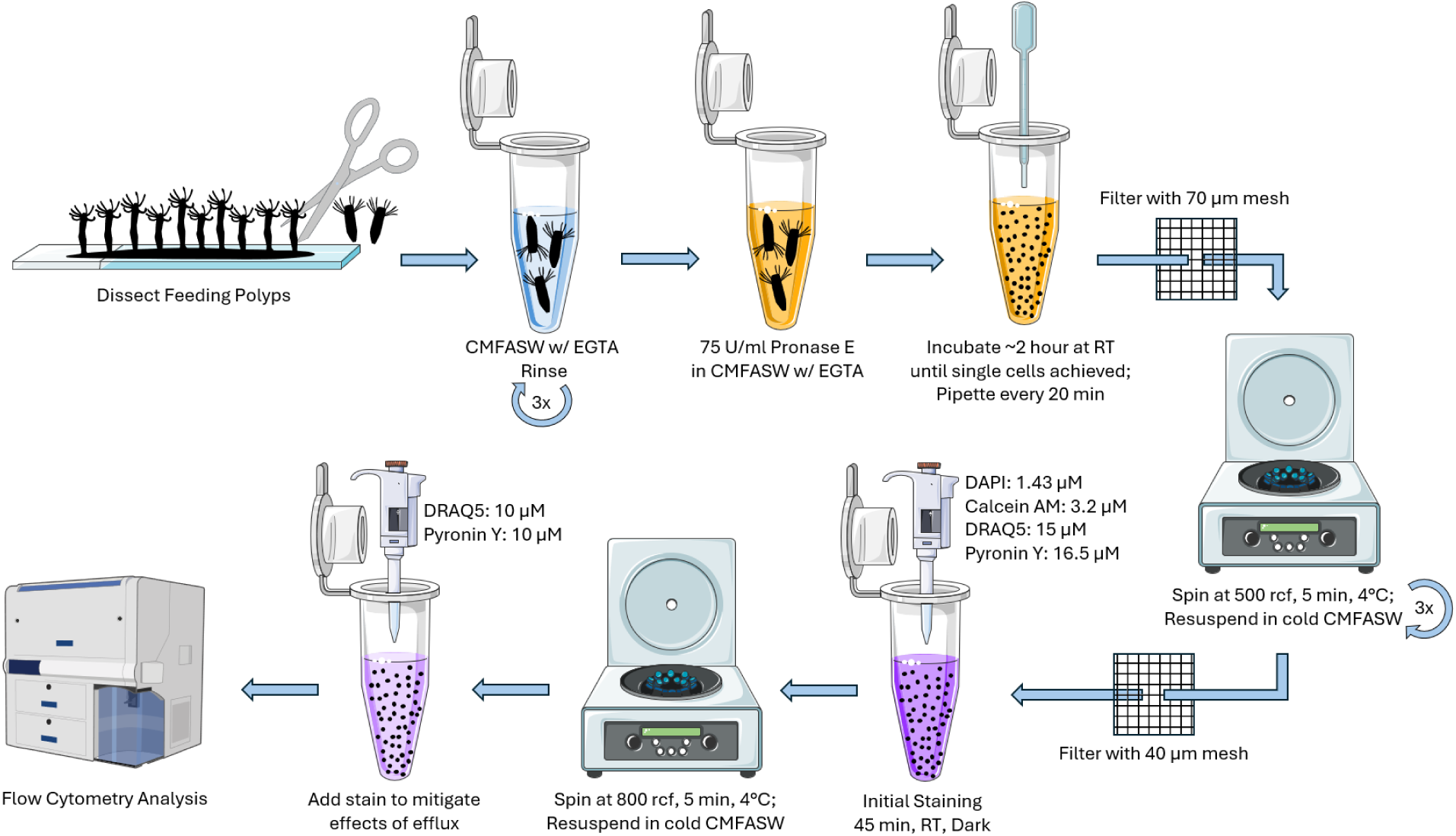
Visual summary of cell dissociation protocol and staining technique for flow cytometry analysis.

**Figure S2.**
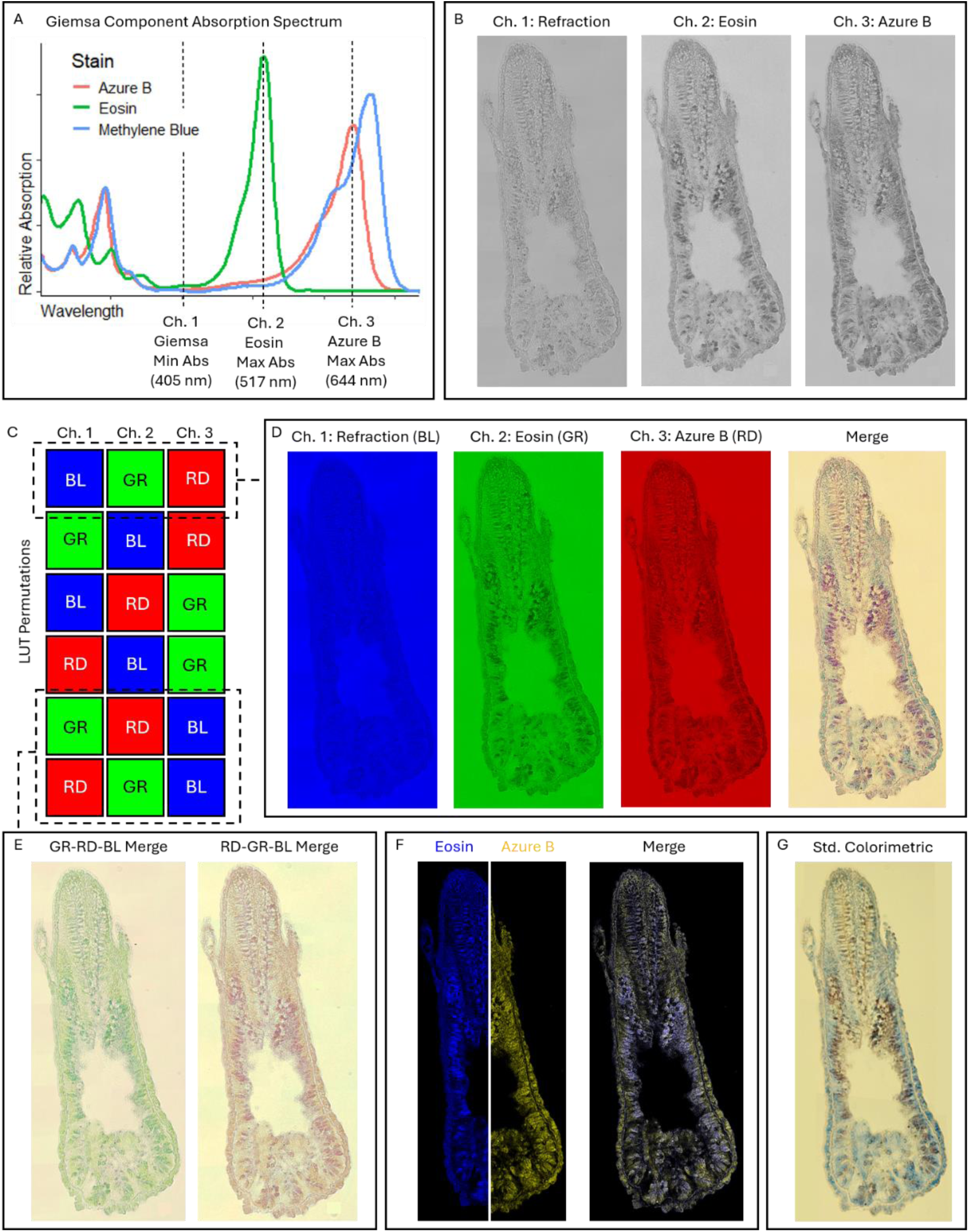
Demonstration of color transmitted light imaging technique. (A) Absorption spectrum of Giemsa components with laser wavelength of each channel labeled (absorption data from AAT Bioquest). (B) Appearance of imaging channels before colored LUT application. Areas of high stain concentration appear as shadows. (C) All possible LUT permutations using red, blue, and green. (D) The LUT permutation used throughout the current study. (E) Two other LUT permutations to demonstrate the arbitrary nature of final image coloration. (F) Use of channel inversion to achieve LUT coloration positively related to stain concentration. (G) Standard colorimetric image for comparison.

**Figure S3.**
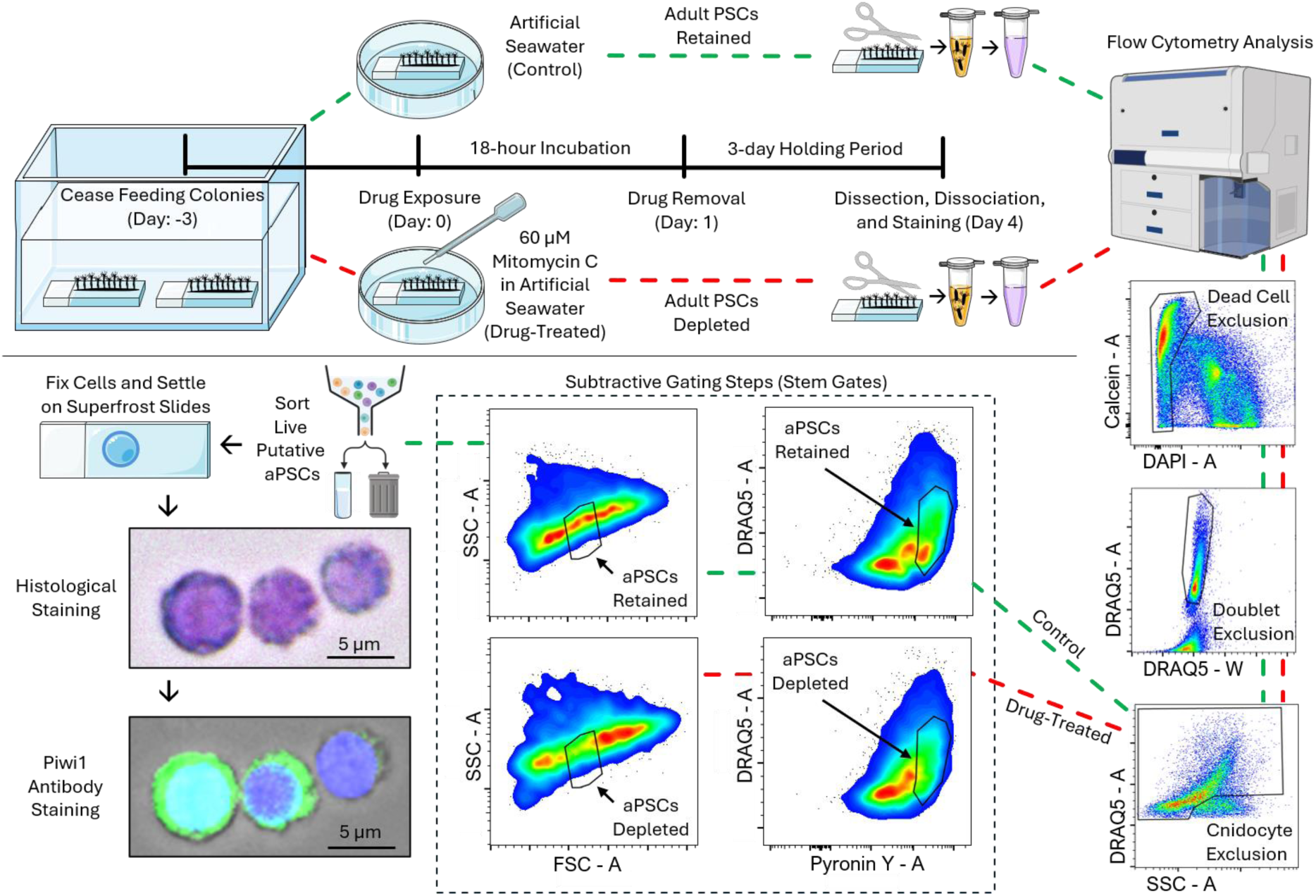
Visual summary of mitomycin experiments, gating strategies, and sorted cell identification approach.

**Figure S4.**
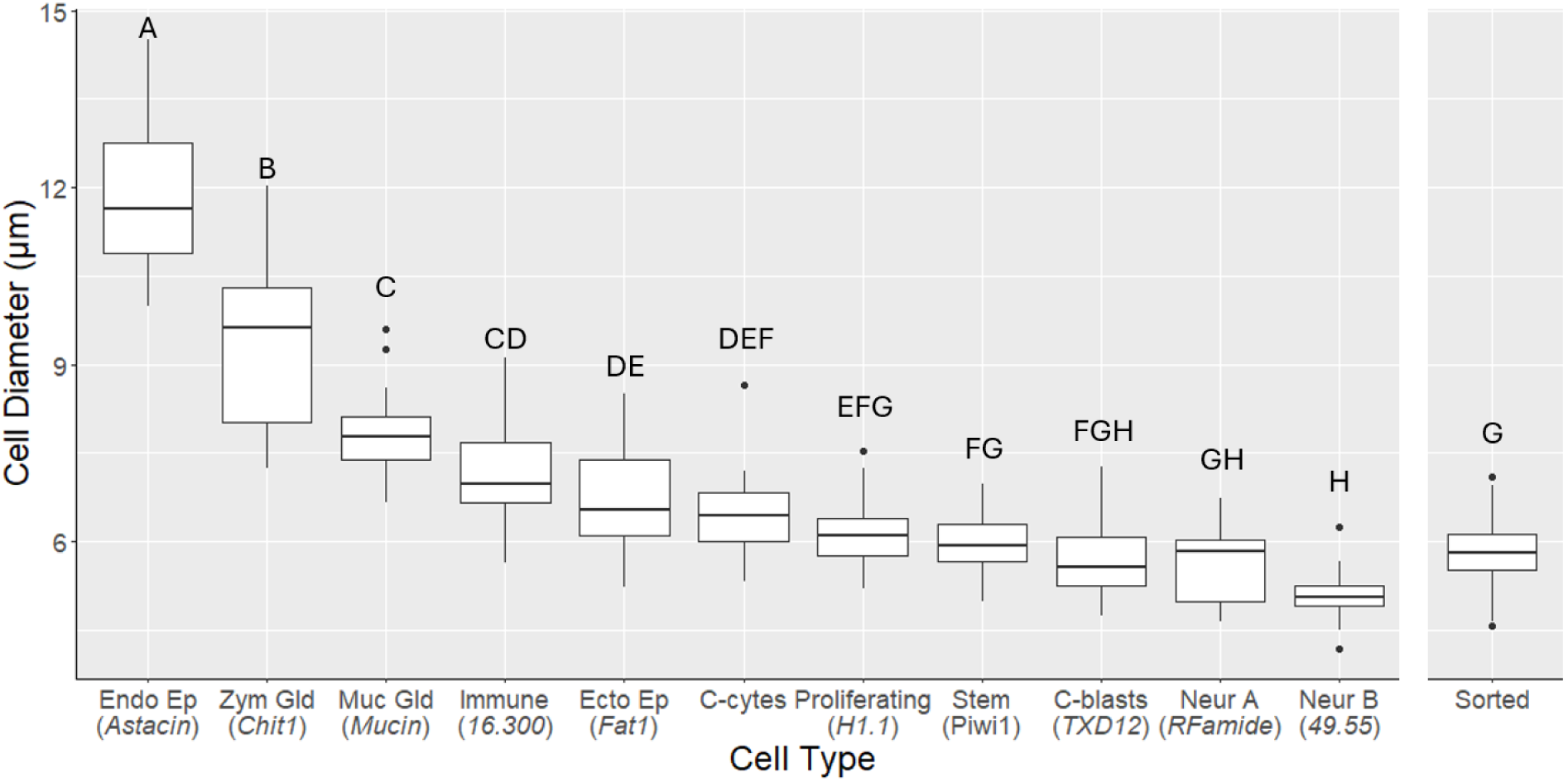
Plot showing differences in cell diameter between cell types. Significant differences according to One-way ANOVA and post-hoc pairwise analysis with α = 0.05 are indicated in standard compact letter display format. Endo Ep = endodermal epithelial; Zym Gld = zymogen gland; Muc Gld = Mucous Gland; Ecto Ep = ectodermal epithelial; C-cytes = cnidocytes; C-blasts = cnidoblasts; Neur = Neuron. Endo Ep n = 10; Sorted n = 100; all others n = 22. *HyS001c.300* and *HyS004S.55* are abbreviated.

**Table S1.**
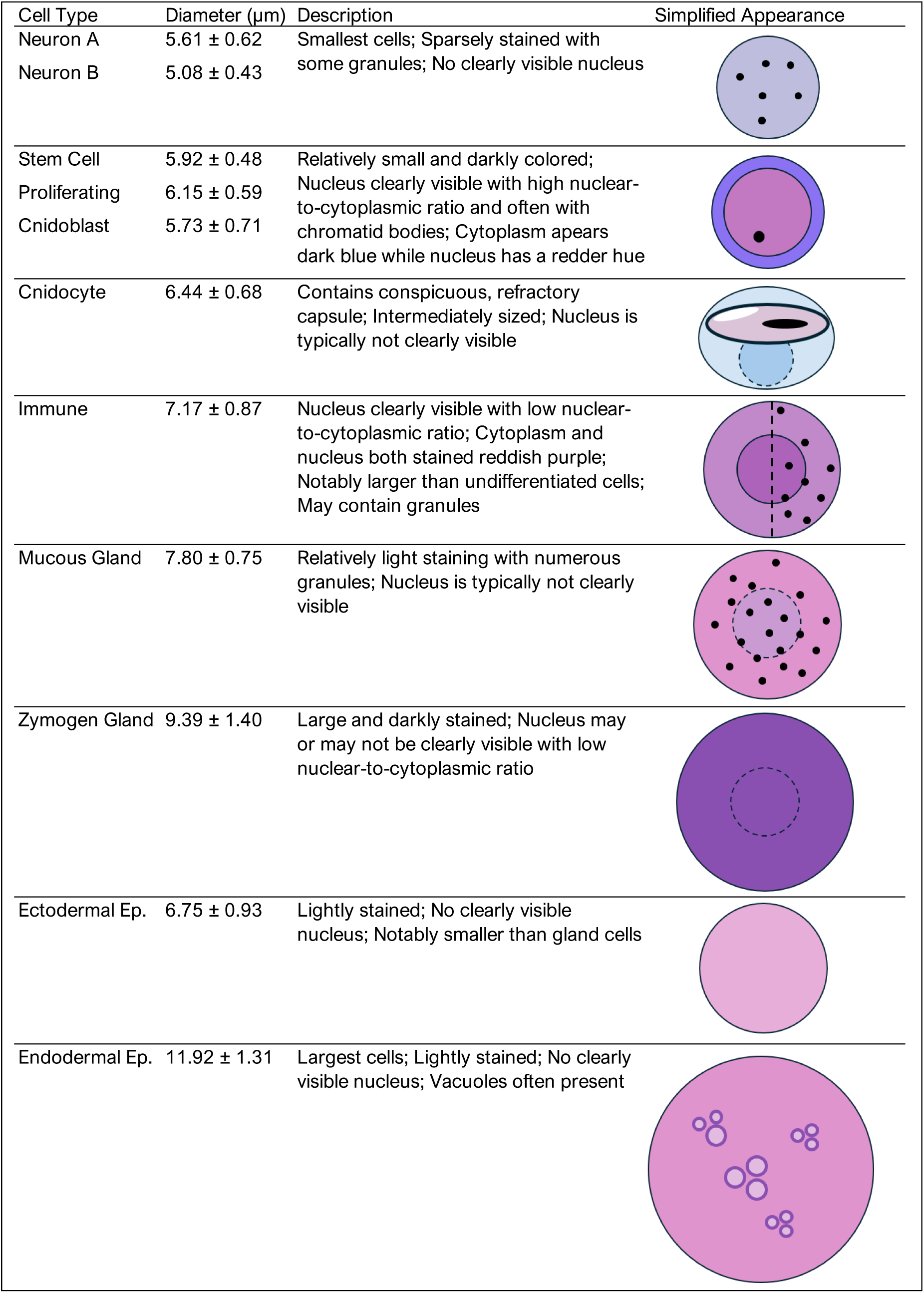
Summary of cytological assay for identifying *Hydractinia* cell types.

| Cell Type | Diameter ($\mu\text{m}$) | Description | Simplified Appearance |
| --- | --- | --- | --- |
| Neuron A<br>Neuron B | $5.61 \pm 0.62$<br>$5.08 \pm 0.43$ | Smallest cells; Sparsely stained with some granules; No clearly visible nucleus | |
| Stem Cell<br>Proliferating<br>Cnidoblast | $5.92 \pm 0.48$<br>$6.15 \pm 0.59$<br>$5.73 \pm 0.71$ | Relatively small and darkly colored; Nucleus clearly visible with high nuclear-to-cytoplasmic ratio and often with chromatid bodies; Cytoplasm appears dark blue while nucleus has a redder hue | |
| Cnidocyte | $6.44 \pm 0.68$ | Contains conspicuous, refractory capsule; Intermediately sized; Nucleus is typically not clearly visible | |
| Immune | $7.17 \pm 0.87$ | Nucleus clearly visible with low nuclear-to-cytoplasmic ratio; Cytoplasm and nucleus both stained reddish purple; Notably larger than undifferentiated cells; May contain granules | |
| Mucous Gland | $7.80 \pm 0.75$ | Relatively light staining with numerous granules; Nucleus is typically not clearly visible | |
| Zymogen Gland | $9.39 \pm 1.40$ | Large and darkly stained; Nucleus may or may not be clearly visible with low nuclear-to-cytoplasmic ratio | |
| Ectodermal Ep. | $6.75 \pm 0.93$ | Lightly stained; No clearly visible nucleus; Notably smaller than gland cells | |
| Endodermal Ep. | $11.92 \pm 1.31$ | Largest cells; Lightly stained; No clearly visible nucleus; Vacuoles often present | |

